# Deconvolution-derived cell-type expression targets for personal genome sequence-to-expression prediction

**DOI:** 10.64898/2026.07.20.739672

**Authors:** Stephen Sim, Li Shen

## Abstract

Sequence-to-function models learn regulatory features from genomic sequence, but they remain limited in their ability to predict gene-expression differences among individuals. Cell-type-specific regulatory effects may be obscured in bulk RNA sequencing, whereas paired genotype and single-cell expression cohorts remain small. We evaluated whether deconvolution of bulk RNA-seq could provide scalable cell-type-specific targets for personal-genome expression prediction. GTEx v8 bulk RNA-seq from six tissues was deconvolved with BayesPrism using single-nucleus reference profiles, producing targets across 83 tissue–cell-type contexts. Deconvolved expression agreed with matched pseudobulked GTEx single-nucleus RNA-seq, with median donor-level Pearson correlations across genes ranging from 0.53 to 0.73 by tissue. We compared genotype-feature models, regressors trained on frozen Enformer representations, and fine-tuned Enformer and Borzoi models. Across random and nonlinear-enriched gene sets, sequence-derived approaches generally outperformed genotype-feature baselines, while frozen Enformer features were competitive with end-to-end fine-tuning. For the random gene set, Fisher-averaged Pearson correlations were 0.122–0.142 for sequence-derived approaches and 0.081–0.086 for genotype-feature baselines in a coverage-aware sensitivity analysis. Model performance was positively associated with deconvolution–pseudobulk agreement for sequence-derived models (*r* = 0.35–0.43 across tissue–cell-type contexts), suggesting that target reliability may constrain downstream prediction. Context-specific Enformer fine-tuning did not materially out-perform a shared, combined-context strategy. These results support deconvolution as a feasible approach for generating cell-type-resolved training targets, while showing that target quality and limited cohort size remain important constraints. Frozen pretrained representations provide a computationally efficient and competitive baseline for personal sequence-to-expression modeling.

## 1 Introduction

Sequence-to-function models have improved the prediction of gene-regulatory activity from genomic sequence. Enformer and Borzoi, for example, combine convolutional and attention-based architectures to predict thousands of genomic tracks from long DNA sequences [1, 2]. These models capture regulatory sequence features across genes and genomic contexts, but success in predicting average regulatory activity does not necessarily translate into prediction of expression differences among individuals. Evaluations using paired whole-genome sequencing (WGS) and RNA-seq have shown that current models often explain little cross-individual expression variation and may fail to predict the direction of cis-regulatory effects [3, 4].

One potential limitation is the use of bulk RNA-seq targets. A bulk measurement averages expression across heterogeneous cell populations, potentially obscuring genetic effects that operate within particular cell types. Cell-type-aware quantitative-trait-locus studies have identified regulatory associations that are weak or undetectable in bulk tissue [5, 6]. Cell-type-resolved expression may therefore provide more informative targets for models intended to predict the functional consequences of personal genetic variation.

Population-scale single-cell and single-nucleus RNA-seq resources remain much smaller than bulk-expression cohorts, particularly when paired genotypes are required. GTEx v8 contains 15,201 bulk RNA-seq samples across 49 tissues from 838 donors, whereas the GTEx cross-tissue singlenucleus atlas includes 25 samples from 16 donors across eight tissues [7, 8]. Cell deconvolution offers a possible bridge between these resources: single-cell or single-nucleus profiles define reference cell types, and large bulk cohorts provide donor-level observations from which cell-type-specific expression can be estimated.

Here, we tested whether deconvolution-derived cell-type expression estimates can serve as targets for personal genome sequence-to-expression prediction (Figure 1). We first assessed agreement between BayesPrism estimates and matched pseudobulked single-nucleus expression. We then compared three modeling strategies: conventional genotype-feature baselines, regressors trained on frozen Enformer representations, and end-to-end fine-tuning of Enformer and Borzoi. Finally, we examined whether variation in deconvolution agreement was associated with downstream prediction performance and whether combined-context fine-tuning improved over context-specific models. We hypothesized that sequence-derived representations would outperform genotype-feature baselines, but that target reliability would limit the benefit of end-to-end fine-tuning.

**Figure 1:**
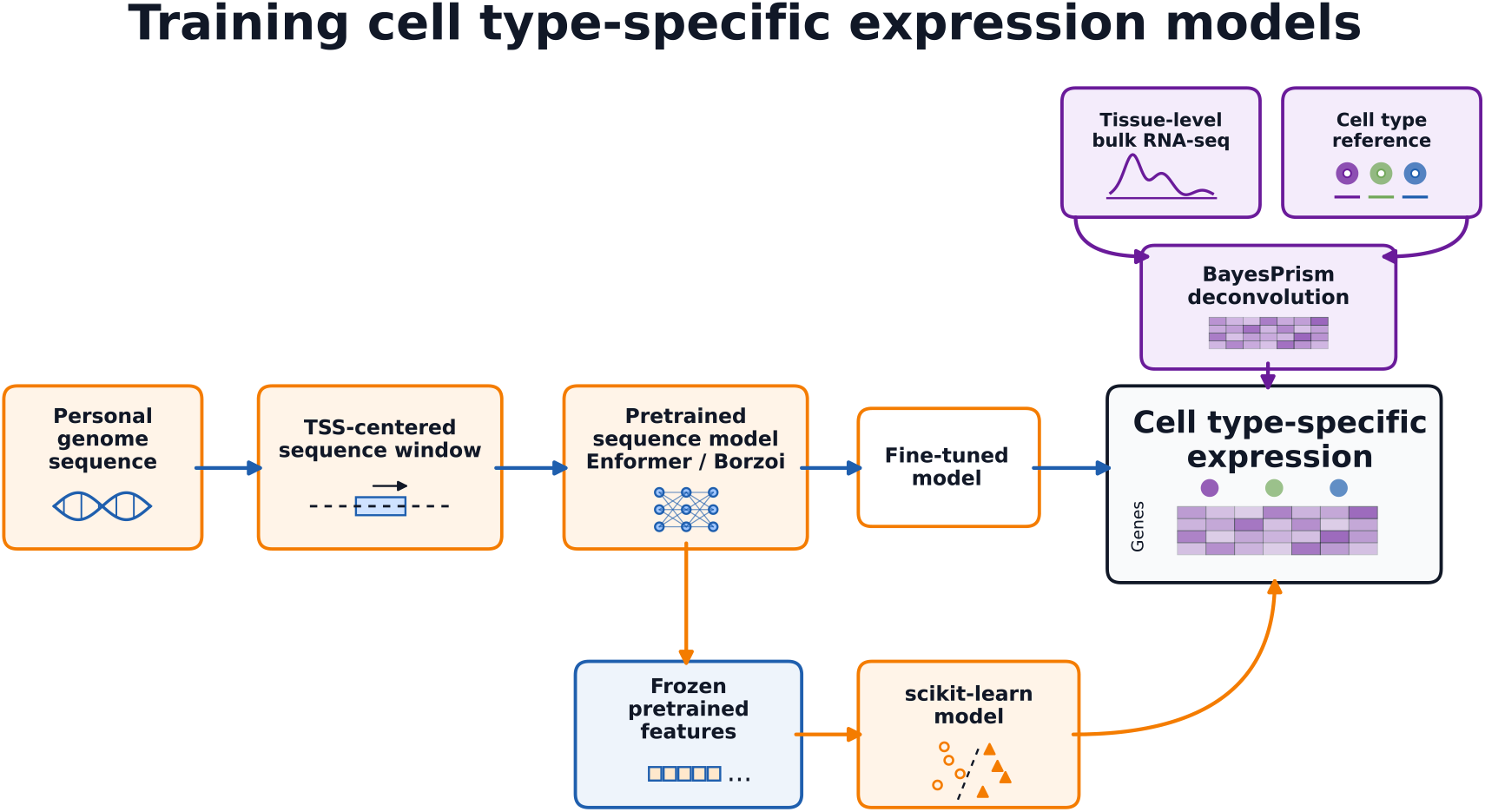
Study overview. Bulk GTEx RNA-seq was deconvolved using single-nucleus reference profiles to generate donor-level cell-type expression targets. Personalized sequence or genotype-derived features were used to predict cross-individual expression. We compared genotype-feature baselines, regressors trained on frozen Enformer representations, and fine-tuned Enformer and Borzoi models.

## 2 Results

### 2.1 Deconvolution produced cell-type-specific targets with variable agreement across tissues

BayesPrism estimates were compared with pseudobulked GTEx single-nucleus RNA-seq for donors with matched data. Pearson correlation was calculated across genes for each donor–tissue–celltype combination. Deconvolved and pseudobulked profiles were positively correlated across all six evaluated tissues (Figure 2). Median donor-level correlations ranged from 0.532 in heart left ventricle to 0.727 in skin (Table 1). The spread within tissues indicated substantial variation among cell types and donors.

**Figure 2:**
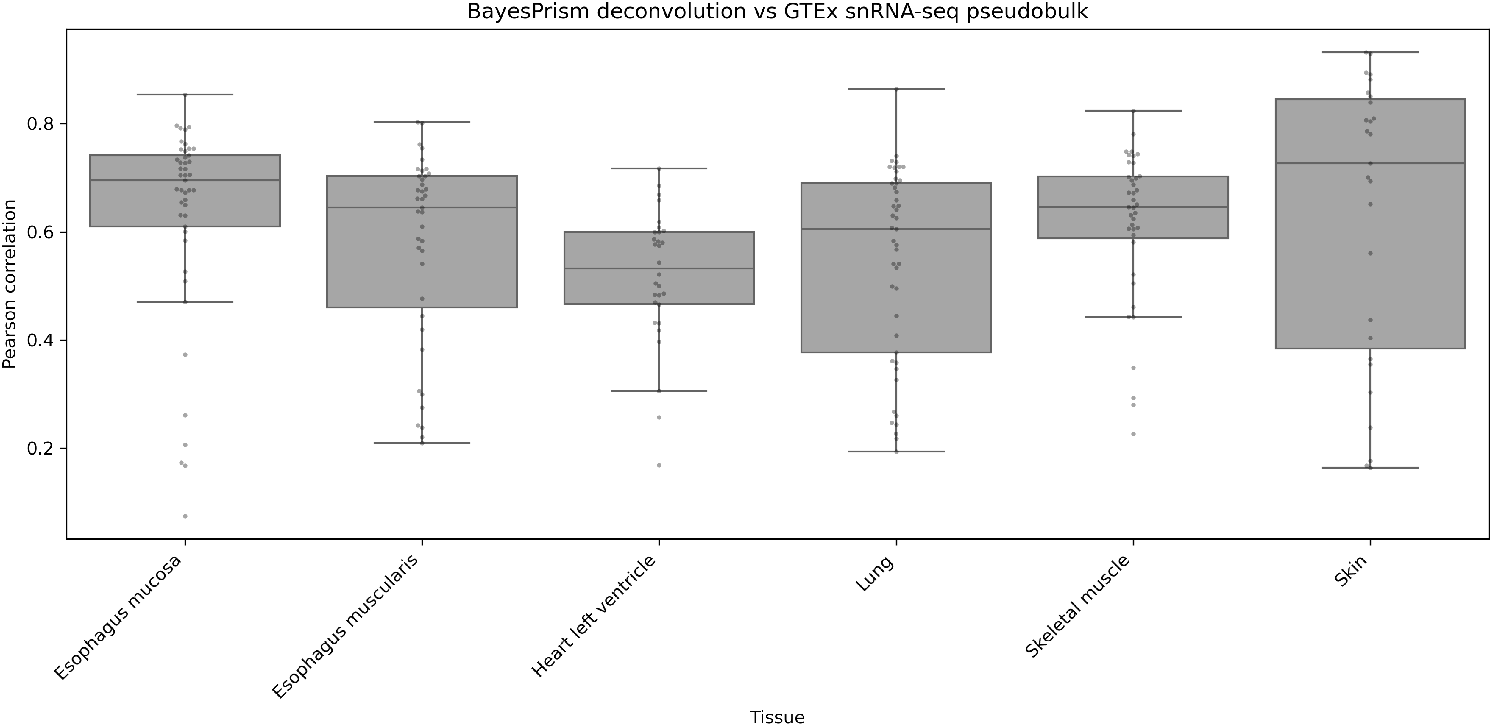
Agreement between BayesPrism-deconvolved expression and pseudobulked GTEx single-nucleus RNA-seq. Each point represents a donor–tissue–cell-type combination, with Pearson *r* calculated across genes. Boxplots summarize the distribution within each tissue.

**Table 1:** Median deconvolution–pseudobulk agreement by tissue.

| Tissue | Median Pearson $r$ |
| --- | --- |
| Esophagus mucosa | 0.695 |
| Esophagus muscularis | 0.645 |
| Heart, left ventricle | 0.532 |
| Lung | 0.605 |
| Skeletal muscle | 0.646 |
| Skin, sun-exposed lower leg | 0.727 |

These comparisons support the use of deconvolution-derived estimates as approximate cell-type-specific targets, but they do not establish equivalence to directly measured cell-type expression. In particular, reference composition, cell-type annotation, and differences between nuclear and whole-cell RNA may contribute to context-dependent agreement.

### 2.2 Sequence-derived representations outperformed genotype-feature baselines

We evaluated model performance across gene–tissue–cell-type contexts in a randomly sampled gene set and a nonlinear-enriched gene set. Because some contexts yielded undefined correlations, we report both matched-context analyses and a coverage-aware sensitivity analysis in which undefined correlations were assigned *r* = 0. The latter jointly reflects prediction accuracy and the proportion of contexts for which a model produced variable predictions, but it should not be interpreted as an unbiased estimate of correlation.

In the random gene set, sequence-derived approaches clustered within a narrow range of Fisher-averaged correlations (0.122–0.142) in the coverage-aware analysis, whereas genotype-feature baselines achieved 0.081–0.086 (Table 2; Figure 3). The nonlinear-enriched set showed the same broad ranking, with sequence-derived models at 0.132–0.170 and genotype-feature models at 0.021–0.096. Because the nonlinear-enriched genes were selected using the relative performance of Random Forest and Elastic Net genotype models, we regard this gene set as exploratory and emphasize the random-gene results.

**Table 2:** Coverage-aware model performance. Values are Fisher-averaged Pearson correlations after undefined context-level correlations were assigned *r* = 0.

| Model | Random genes | Nonlinear-enriched genes |
| --- | --- | --- |
| Borzoi, fine-tuned | 0.139 | 0.145 |
| Enformer, fine-tuned | 0.134 | 0.133 |
| Enformer, contrastive loss | 0.142 | 0.144 |
| Enformer, unphased input | 0.133 | 0.132 |
| Elastic Net, Enformer embeddings | 0.141 | 0.170 |
| Random Forest, Enformer embeddings | 0.130 | 0.152 |
| Elastic Net, Enformer output tracks | 0.133 | 0.158 |
| Random Forest, Enformer output tracks | 0.122 | 0.140 |
| Elastic Net, genotype features | 0.081 | 0.021 |
| Random Forest, genotype features | 0.086 | 0.096 |

**Figure 3:**
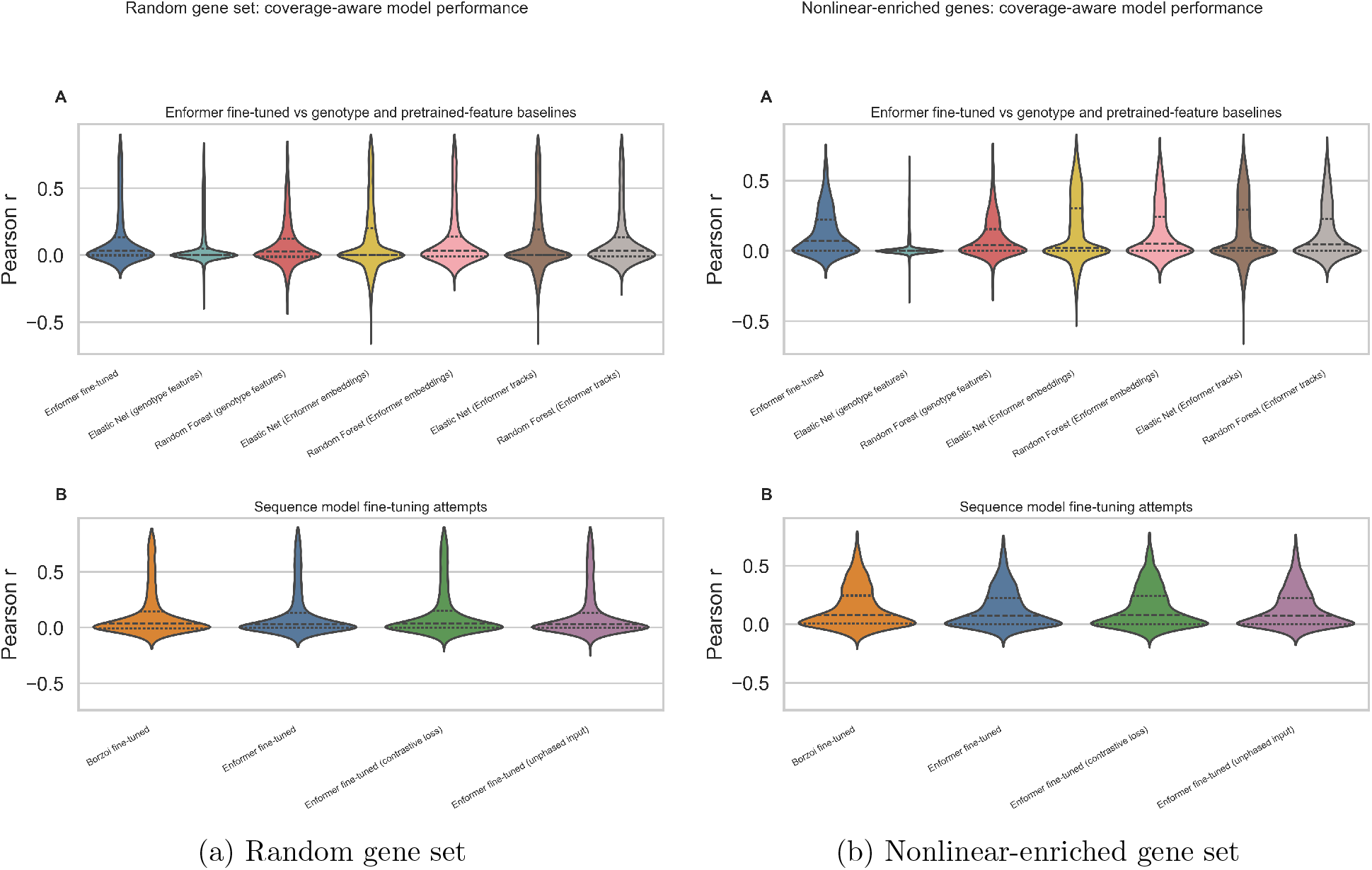
Coverage-aware cross-individual prediction performance. Distributions show context-level Pearson correlations after undefined values were assigned *r* = 0. This analysis is presented as a sensitivity analysis; matched-context results and coverage should be considered alongside it.

### 2.3 Frozen pretrained representations were competitive with end-to-end finetuning

Fine-tuned Enformer and Borzoi models did not consistently outperform regressors trained on frozen Enformer representations. Elastic Net using Enformer embeddings achieved the highest coverage-aware Fisher-averaged correlation in the nonlinear-enriched set, while Enformer with contrastive loss was highest in the random set; however, differences among sequence-derived methods were small relative to their context-level variability.

Pairwise comparisons showed that the globally similar averages concealed large differences for individual gene–tissue–cell-type contexts (Supplementary Figure 5). Some contexts favored Borzoi fine-tuning, whereas others favored Elastic Net on frozen Enformer embeddings. Thus, no sequence-derived strategy was uniformly superior. The competitiveness of frozen features is notable because they require only pretrained-model inference followed by conventional regression, avoiding repeated end-to-end GPU fine-tuning.

### 2.4 Deconvolution agreement was associated with downstream model performance

Across tissue–cell-type contexts, higher deconvolution–pseudobulk agreement was associated with better prediction by sequence-derived models (Figure 4). Cross-context correlations ranged from 0.345 to 0.427 for fine-tuned models and regressors trained on frozen Enformer features (Table 3). In contrast, genotype-feature baselines showed near-zero or negative associations (*r* = −0.107 and −0.034).

**Figure 4:**
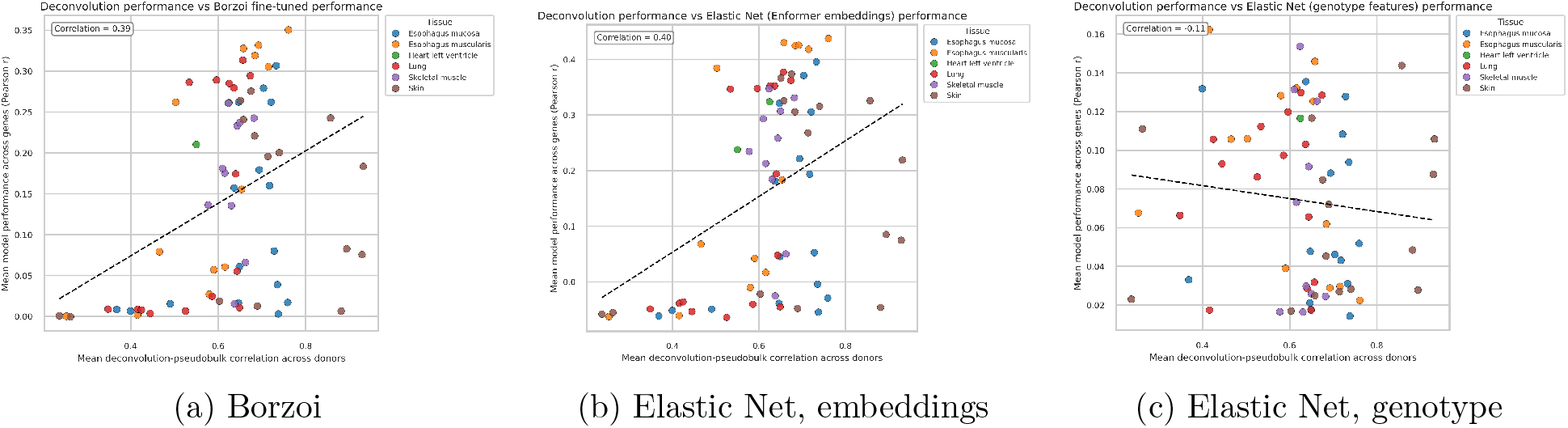
Association between deconvolution agreement and model performance. Each point is a tissue–cell-type context. The horizontal axis is mean BayesPrism–pseudobulk Pearson correlation across matched donors; the vertical axis is mean prediction Pearson *r* across genes.

**Table 3:** Cross-context association between deconvolution agreement and prediction performance.

| Model | Pearson $r$ |
| --- | --- |
| Enformer, fine-tuned | 0.427 |
| Enformer, contrastive loss | 0.425 |
| Enformer, unphased input | 0.423 |
| Elastic Net, Enformer output tracks | 0.403 |
| Elastic Net, Enformer embeddings | 0.401 |
| Borzoi, fine-tuned | 0.388 |
| Random Forest, Enformer embeddings | 0.346 |
| Random Forest, Enformer output tracks | 0.345 |
| Random Forest, genotype features | -0.034 |
| Elastic Net, genotype features | -0.107 |

This analysis is associative. Tissue identity, cell-type composition, sample size, expression variability, and reference quality could influence both deconvolution agreement and prediction performance. Nevertheless, the consistency across sequence-derived approaches suggests that target reliability may be an important constraint on cross-individual prediction.

### 2.5 Context-specific fine-tuning provided no material advantage

For 25 genes from the nonlinear-enriched set, we compared combined-context Enformer models, which shared a fine-tuned backbone across tissue targets, with models trained separately for individual tissue–cell-type contexts. Among 2,321 contexts with valid correlations for both strategies, model performance was highly concordant (*r* = 0.902). Mean Pearson correlation was 0.149 for combined-context models and 0.157 for context-specific models, a paired mean difference of 0.0085. A sensitivity analysis assigning undefined correlations to zero produced the same conclusion across 2,520 contexts (means 0.137 and 0.151; cross-model *r* = 0.873; Supplementary Figure 6). Combined-context fine-tuning therefore did not provide an evident performance advantage in this experiment.

## 3 Discussion

We evaluated deconvolution-derived cell-type expression as a scalable target for predicting expression variation from personal genomes. Three findings emerged. First, BayesPrism estimates showed moderate-to-strong agreement with pseudobulked single-nucleus profiles, but agreement varied across tissues and cell types. Second, sequence-derived approaches generally outperformed models trained directly on local genotype features. Third, frozen Enformer representations were competitive with end-to-end fine-tuning, and downstream performance was associated with deconvolution agreement.

The deconvolution results support feasibility rather than equivalence to direct single-cell measurement. Reference profiles were derived from nuclei, whereas bulk RNA-seq captures nuclear and cytoplasmic RNA. Reference composition, annotation granularity, and the small number of matched GTEx single-nucleus donors can affect the comparison. Deconvolution benchmarks based on pseudobulk data may also fail to reproduce the biological variability of genuine bulk tissue [9]. Frozen Enformer features provided a strong computationally efficient baseline. Pretrained representations may already encode regulatory features useful for cross-individual prediction, allowing a regularized downstream model to extract relevant variation without updating millions of deep-model parameters. This interpretation is consistent with prior work using pretrained sequence embeddings for personal expression prediction [10]. Conversely, end-to-end fine-tuning may be particularly difficult when donor counts are modest and labels contain deconvolution error. Large neural networks can fit noisy targets, and stronger regularization or lower-capacity adaptation may generalize better in this regime [11, 12].

Our comparisons also emphasize that average performance can hide considerable context dependence. Fine-tuned models and frozen-feature regressors performed similarly in aggregate but differed substantially for particular genes, tissues, and cell types. Identifying the characteristics that favor each modeling strategy may be more informative than seeking a single global winner. The absence of a clear advantage for combined-context fine-tuning further suggests that sharing information across the evaluated contexts was insufficient to overcome target noise or limited sample size.

Several limitations qualify these findings. The nonlinear-enriched gene set was selected using baseline-model performance and is therefore susceptible to selection bias; the random-gene analysis is the more appropriate basis for general conclusions. Assigning undefined correlations to zero conflates coverage and accuracy, so this analysis should remain secondary to paired matched-context comparisons with coverage reported separately. Finally, contexts are statistically dependent because they share genes, donors, tissues, and cell types.

Future work should improve both target generation and model adaptation. Leave-one-donor-out deconvolution, comparisons across reference atlases and deconvolution methods, and target-reliability weights could clarify how label quality affects training. Adapter layers, partial fine-tuning, or low-rank updates may offer a middle ground between frozen representations and full fine-tuning. Larger paired genotype and cell-type-resolved expression cohorts will ultimately be necessary to determine whether sequence models can robustly predict cell-type-specific expression variation among individuals.

In summary, deconvolution can expand the scale of cell-type-resolved expression targets available for personal-genome modeling, but target reliability remains a central constraint. Pretrained sequence representations were more informative than direct genotype-feature baselines and performed comparably to end-to-end fine-tuning at substantially lower computational cost. These findings establish frozen representations as an important baseline and motivate more rigorous evaluation of deconvolution-derived targets.

## 4 Methods

### 4.1 Datasets and personal sequence construction

Bulk RNA-seq and genotype data were obtained from GTEx v8. GTEx single-nucleus data were obtained from the cross-tissue reference atlas; Heart Cell Atlas data provided the cardiac reference; and ROSMAP genotype and single-nucleus data were used for the brain reference and external evaluation [7, 8, 13, 14]. The main comparative analyses used esophagus mucosa, esophagus muscularis, heart left ventricle, lung, skin, and skeletal muscle. Brain analyses were used during resource development and are described in the Supplementary Methods.

Personalized inputs were defined in 49,152-bp windows centered on each transcription start site, matching prior work [15]. The hg38 genome was used as reference. Variants were parsed with cyvcf2, and biallelic SNPs and indels were applied with bcftools [16, 17]. Sequences were adjusted after indel application to preserve fixed-length, TSS-centered windows and were one-hot encoded with kipoiseq [18, 19].

### 4.2 Cell-type deconvolution and target processing

GTEx bulk RNA-seq was deconvolved with BayesPrism, which estimates cell proportions and cell-type-specific expression from a single-cell or single-nucleus reference [20]. GTEx single-nucleus references were used for most tissues. Heart Cell Atlas nuclei from left ventricle were used for the cardiac reference, and healthy ROSMAP nuclei were used for frontal cortex. Reference matrices were downsampled to 200 cells per cell type. Protein-coding genes were retained after excluding sex-chromosome, ribosomal, and mitochondrial genes according to the BayesPrism workflow.

Cell-type-specific matrices were normalized by the trimmed mean of M-values method in edgeR and transformed with voom in limma. Tissue-specific covariates were regressed from expression; each gene’s mean log-counts-per-million was added back to retain its expression scale [21, 22]. GTEx single-nucleus counts were pseudobulked by donor, tissue, and cell type with decoupler and processed using the same normalization and residualization procedure [23].

### 4.3 Genotype-feature baseline models

Elastic Net and Random Forest baselines were implemented in `scikit-learn` [24]. For each gene, phased variants within the 49,152-bp TSS-centered window were encoded as concatenated haplotype indicators. Features with training-set alternate-allele frequency below 0.01 were removed, and missing genotype values were imputed using training-set means. MultiTaskElasticNetCV used default hyperparameters and `random_state=8888`; RandomForestRegressor used 300 trees, `n_jobs=-1`, and `random_state=8888`. Models were evaluated with shuffled five-fold cross-validation. Missing multitask targets were imputed within each fold using MissForest fit on the training matrix [25].

### 4.4 Gene-set selection

Two gene sets were evaluated. A nonlinear-enriched set was identified by screening contexts in which Random Forest genotype models outperformed Elastic Net genotype models. Bootstrap confidence intervals (*n* = 1,000) were converted to approximate standard errors, and two-sided *P* values for the difference in Pearson correlation were adjusted using the Benjamini–Hochberg procedure [26]. This produced 144 unique genes and was treated as an exploratory, selection-enriched set.

For the random set, context-level baseline correlations were divided into ten equal-width bins, and 30 contexts were sampled per bin. Because sampling was performed over contexts, duplicate genes were possible; 300 sampled contexts corresponded to 292 unique genes. Overlap with the nonlinear-enriched set was allowed.

### 4.5 Models using frozen Enformer representations

For each 49,152-bp sequence, the two haplotypes of each individual were independently processed using Enformer with frozen parameters. For each haplotype, Enformer generated position-by-feature tensors of dimensions *L ×* 3,072 and *L ×* 5,313, corresponding to the Enformer embedding channels and human output-head tracks, respectively. At Enformer’s 128-bp resolution, *L* = 384 positional bins. For each tissue *t*, the 5,313 output tracks were filtered to retain only a predefined subset of *T*_*t*_ tissue-relevant tracks, yielding a tensor of dimensions *L × T*_*t*_. Both representations were then mean-pooled across the *L* positional bins, producing a 3,072-dimensional embedding vector and a *T*_*t*_-dimensional output-track vector for each haplotype. Finally, the corresponding vectors from the two haplotypes were averaged elementwise, yielding one embedding vector for each gene–individual pair and one tissue-specific output-track vector for each gene–tissue–individual combination.

Elastic Net and Random Forest were trained separately for each gene–tissue–cell-type context. ElasticNetCV used *ℓ*_1_ ratios of 0.01, 0.5, and 0.99 and alpha values from 10^−3^ to 10^2^. RandomForestRegressor used 100 trees, unrestricted depth, minimum leaf size 1, and `random_state=88`. Features were standardized within each training fold. Models were evaluated with shuffled five-fold cross-validation.

### 4.6 Fine-tuning Enformer and Borzoi

Enformer and Borzoi were implemented in PyTorch using `enformer-pytorch` and `borzoi-pytorch`, respectively [27–30]. Each model received 49,152-bp personalized sequences. Haplotype inputs were processed separately, and the two predictions were averaged to produce the final output; unphased inputs were evaluated as a sensitivity analysis. Gene-specific models used tissue-specific output heads and a shared backbone across available contexts. Borzoi embeddings were connected to one-dimensional convolutional output heads.

Models were trained with five-fold cross-validation for 30 epochs using batch size 8. Within each training fold, 10% of samples were reserved for validation. AdamW optimization used learning rate 3 *×* 10^−5^, weight decay 0.01, and no dropout. The primary objective was masked mean squared error over observed targets. Contrastive-loss experiments added a pairwise expression-difference objective [15]. The best epoch was selected by Fisher-averaged validation correlation and evaluated on the held-out fold. Each model was trained on one NVIDIA H100 80-GB GPU. Context-specific Enformer models were additionally trained for 25 nonlinear-enriched genes using the same procedure but predicting one gene–tissue–cell-type context per model.

### 4.7 Evaluation

Pearson correlation across individuals was the primary endpoint. Correlations were aggregated by applying Fisher’s *z* = arctanh(*r*) transformation, averaging on the *z* scale, and back-transforming. Undefined correlations were retained as missing in primary paired comparisons. A coverage-aware sensitivity analysis assigned undefined correlations *r* = 0 to represent no demonstrated correlation signal, while recognizing that this combines coverage and accuracy. Out-of-fold predictions were used for evaluation.

## 5 Data and code availability

GTEx data are available through the GTEx Portal and controlled-access genotype and sequencing data through dbGaP under the applicable GTEx accession. ROSMAP data are available through the relevant controlled-access repository. Heart Cell Atlas data are available from the cited resource. Analysis code and processed outputs that can be shared under the applicable data-use agreements will be made available with the published version of this work. No controlled-access individual-level data will be distributed publicly.

## 6 Ethics and controlled-access data

This study used existing controlled-access genomic and transcriptomic data from the Genotype-Tissue Expression (GTEx) project and the Religious Orders Study and Memory and Aging Project (ROSMAP). Access was authorized through the NIH database of Genotypes and Phenotypes (dbGaP) following review and approval of the investigators’ Data Access Requests by the relevant NIH Data Access Committees. The data were accessed and analyzed in accordance with the approved research-use statements, applicable data-use limitations, and NIH Data Use Certification Agreements. Institutional authorization and records of the approved data access are maintained by the Grants and Contracts Office of the Icahn School of Medicine at Mount Sinai. No new participants were recruited and no new biological specimens were collected for this study. Informed consent and ethical oversight for the original data collection were provided by the respective GTEx and ROSMAP studies, as described in their original publications.

## 7 Author contributions

**Stephen Sim:** Methodology, software, formal analysis, investigation, visualization, and writing— original draft. **Li Shen:** Conceptualization, supervision, methodology, project administration, and writing—review and editing.

## 9 Acknowledgements

This work was supported in part through the Minerva computational and data resources and staff expertise provided by Scientific Computing and Data at the Icahn School of Medicine at Mount Sinai and supported by the Clinical and Translational Science Awards grant UL1TR004419 from the National Center for Advancing Translational Sciences. Research reported in this publication was also supported by the Office of Research Infrastructure of the National Institutes of Health under award S10OD030463. The content is solely the responsibility of the authors and does not necessarily represent the official views of the National Institutes of Health. The authors thank Adam Catto for guidance and technical contributions.

## A Supplementary information

### A.1 Supplementary Methods

Seven tissues were initially deconvolved: brain frontal cortex, esophagus mucosa, esophagus mus-cularis, heart left ventricle, lung, sun-exposed skin, and skeletal muscle. The brain frontal-cortex reference used ROSMAP single-nucleus profiles from healthy individuals and retained cell types represented in at least 95% of donors. A brain-enriched gene set was assembled from the Human Protein Atlas single-nucleus brain resource by mapping ROSMAP cell types to the closest atlas categories [31–33].

### A.2 Supplementary Figures

**Figure 5:**
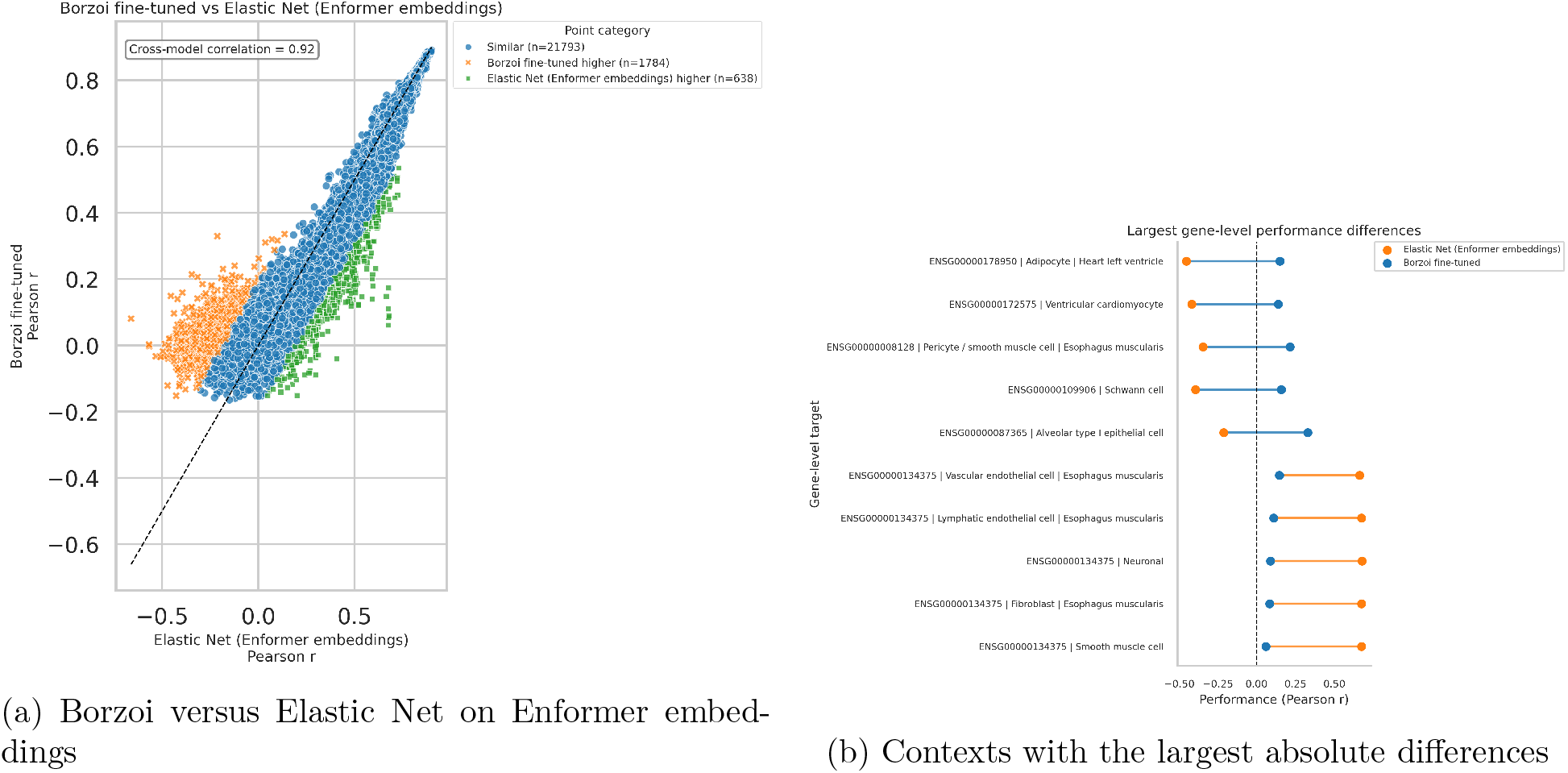
Context-level comparison of Borzoi fine-tuning and Elastic Net trained on frozen Enformer embeddings.

**Figure 6:**
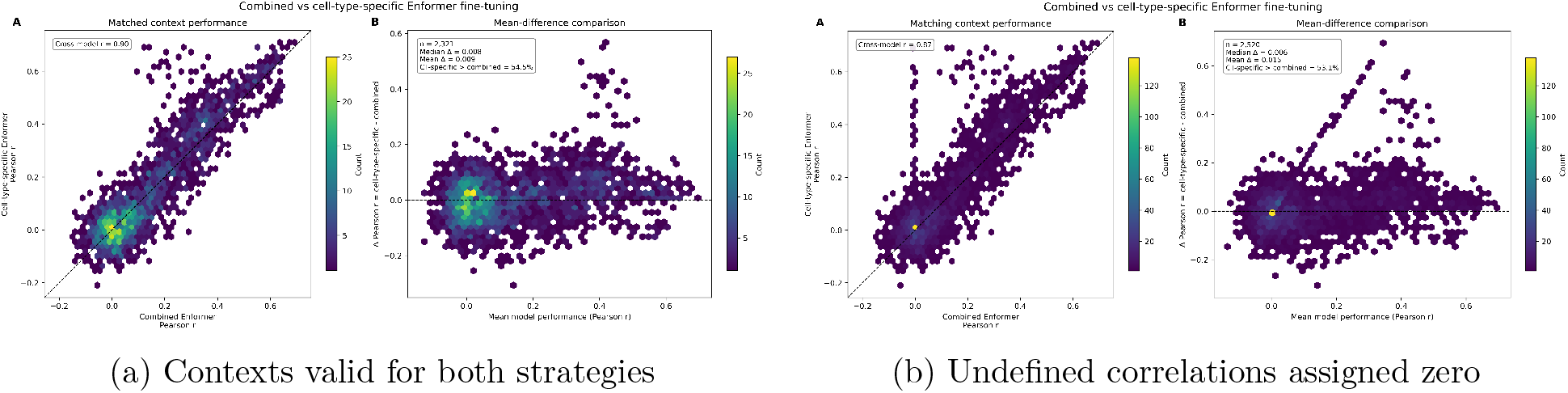
Combined-context versus context-specific Enformer fine-tuning.

**Figure 7:**
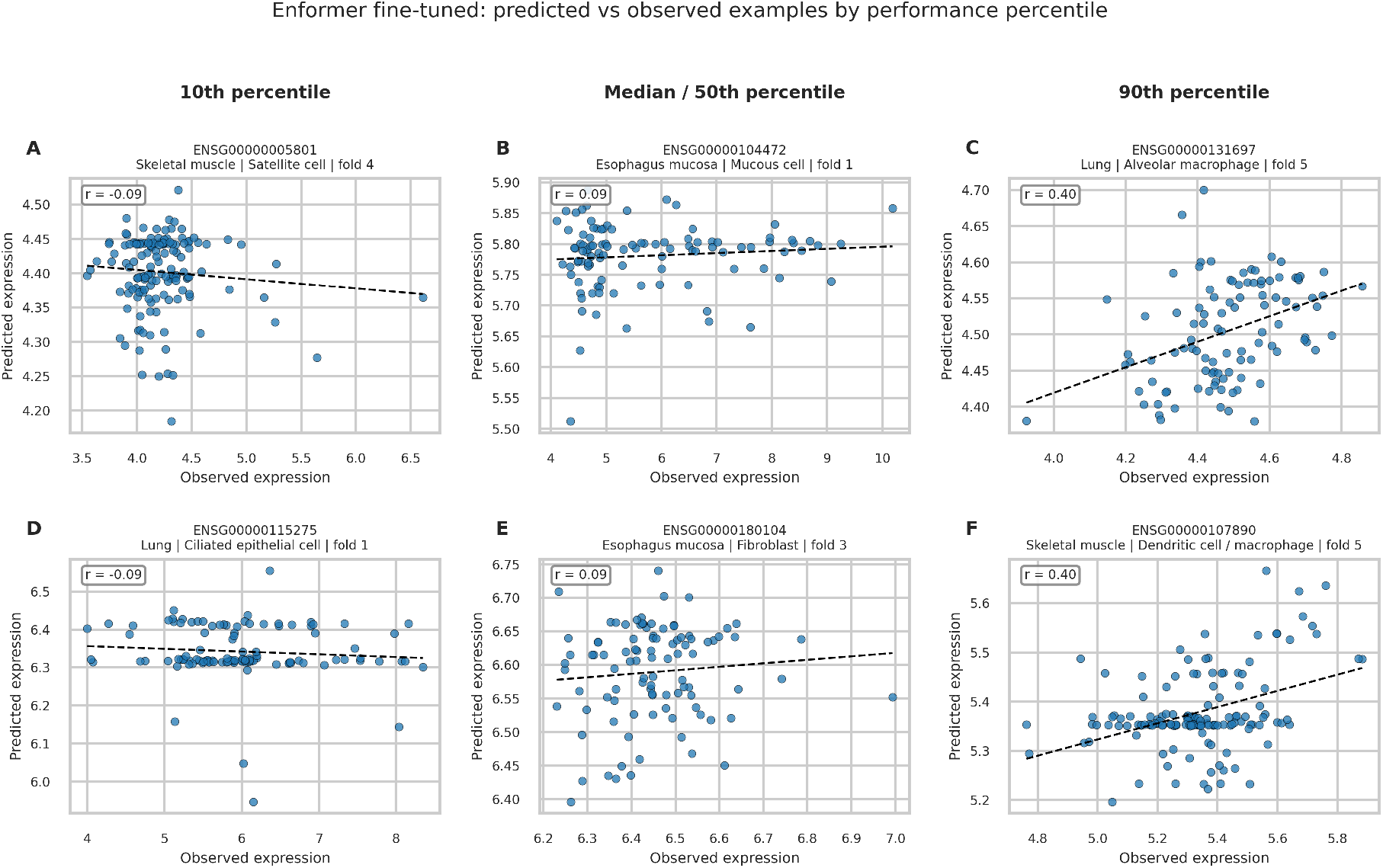
Representative held-out Enformer predictions versus deconvolution-derived targets across gene–tissue–cell-type contexts.

### A.3 Supplementary Tables

**Table 4:** GTEx donors with paired genotype and expression data in the principal tissues.

| Tissue | Paired snRNA-seq-WGS | Paired bulk RNA-seq-WGS |
| --- | --- | --- |
| Esophagus mucosa | 3 | 497 |
| Esophagus muscularis | 3 | 465 |
| Heart, left ventricle | 3 | 386 |
| Lung | 3 | 515 |
| Skin | 3 | 605 |
| Skeletal muscle | 3 | 706 |

**Table 5:** Combined-context and context-specific Enformer fine-tuning.

| Metric | Valid in both models | Undefined $r = 0$ |
| --- | --- | --- |
| Number of contexts | 2,321 | 2,520 |
| Combined-context mean $r$ | 0.149 | 0.137 |
| Context-specific mean $r$ | 0.157 | 0.151 |
| Mean paired difference | 0.0085 | 0.015 |
| Context-specific model better | 54.4% | 53.1% |
| Cross-model correlation | 0.902 | 0.873 |

## References

[1] Žiga Avsec, Vikram Agarwal, Daniel Visentin, Joseph R. Ledsam, Agnieszka Grabska-Barwinska, Kyle R. Taylor, Yannis Assael, John Jumper, Pushmeet Kohli, and David R. Kelley. Effective gene expression prediction from sequence by integrating long-range interactions. Nature Methods, 18(10):1196–1203, 2021. doi: 10.1038/s41592-021-01252-x.

[2] Johannes Linder, Divyanshi Srivastava, Han Yuan, Vikram Agarwal, and David R. Kelley. Predicting RNA-seq coverage from DNA sequence as a unifying model of gene regulation. Nature Genetics, 57(4):949–961, 2025. doi: 10.1038/s41588-024-02053-6.

[3] Connie Huang, Richard W. Shuai, Parth Baokar, Ryan Chung, Ruchir Rastogi, Pooja Kathail, et al. Personal transcriptome variation is poorly explained by current genomic deep learning models. Nature Genetics, 55:2056–2059, 2023. doi: 10.1038/s41588-023-01574-w.

[4] Alexander Sasse, Bernard Ng, Anna E. Spiro, Shinya Tasaki, David A. Bennett, Christopher Gaiteri, Philip L. De Jager, Maria Chikina, and Sara Mostafavi. Benchmarking of deep neural networks for predicting personal gene expression from dna sequence highlights shortcomings. Nature Genetics, 55(12):2060–2064, 2023. doi: 10.1038/s41588-023-01524-6.

[5] Sarah Kim-Hellmuth, François Aguet, Meritxell Oliva, Manuel Muñoz-Aguirre, Silva Kasela, Valentin Wucher, Stephane E. Castel, Andrew R. Hamel, Ana Viñuela, Amy L. Roberts, et al. Cell type-specific genetic regulation of gene expression across human tissues. Science, 369 (6509):eaaz8528, 2020. doi: 10.1126/science.aaz8528.

[6] Christopher D. Brown, Lara M. Mangravite, and Barbara E. Engelhardt. Integrative modeling of eqtls and cis-regulatory elements suggests mechanisms underlying cell type specificity of eqtls. PLOS Genetics, 9(8):e1003649, 2013. doi: 10.1371/journal.pgen.1003649.

[7] GTEx Consortium. The GTEx consortium atlas of genetic regulatory effects across human tissues. Science, 369(6509):1318–1330, 2020. doi: 10.1126/science.aaz1776.

[8] Gökcen Eraslan, Eugene Drokhlyansky, Shankara Anand, Ayshwarya Subramanian, Evgenij Fiskin, Michal Slyper, Jiali Wang, Nicholas Van Wittenberghe, John M. Rouhana, Julia Waldman, Orr Ashenberg, Danielle Dionne, Thet Su Win, Michael S. Cuoco, Olena Kuksenko, Philip A. Branton, Jamie L. Marshall, Anna Greka, Gad Getz, Ayellet V. Segrè, François Aguet, Orit Rozenblatt-Rosen, Kristin G. Ardlie, and Aviv Regev. Single-nucleus cross-tissue molecular reference maps toward understanding disease gene function. Science, 376(6594): eabl4290, 2022. doi: 10.1126/science.abl4290.

[9] Mengying Hu and Maria Chikina. Heterogeneous pseudobulk simulation enables realistic benchmarking of cell-type deconvolution methods. Genome Biology, 25(1):169, 2024. doi: 10.1186/s13059-024-03292-w.

[10] Pratik Ramprasad, Nikhil Pai, and Wei Pan. Enhancing personalized gene expression prediction from DNA sequences using genomic foundation models. Human Genetics and Genomics Advances, 5(4):100347, 2024. doi: 10.1016/j.xhgg.2024.100347.

[11] Hwanjun Song, Minseok Kim, Dongmin Park, Yooju Shin, and Jae-Gil Lee. Learning from noisy labels with deep neural networks: A survey. IEEE Transactions on Neural Networks and Learning Systems, 34(11):8135–8153, 2023. doi: 10.1109/TNNLS.2022.3152527.

[12] Hao Cheng, Zhaowei Zhu, Xing Sun, and Yang Liu. Mitigating memorization of noisy labels via regularization between representations. In International Conference on Learning Representations, 2023. URL https://openreview.net/forum?id=6qcYDVlVLnK.

[13] Alejandra P. Pérez-González, Aidee Lashmi García-Kroepfly, Keila Adonai Pérez-Fuentes, Roberto Isaac García-Reyes, Fryda Fernanda Solis-Roldan, Jennifer Alejandra Alba-González, Enrique Hernández-Lemus, and Guillermo de Anda-Jáuregui. The ROSMAP project: aging and neurodegenerative diseases through omic sciences. Frontiers in Neuroinformatics, 18: 1443865, 2024. doi: 10.3389/fninf.2024.1443865.

[14] Monika Litviňuková, Carlos Talavera-López, Henrike Maatz, Daniel Reichart, Catherine L. Worth, Emma L. Lindberg, et al. Cells of the adult human heart. Nature, 588(7838):466–472, 2020. doi: 10.1038/s41586-020-2797-4.

[15] Shiron Drusinsky, Sean Whalen, and Katherine S. Pollard. Deep-learning prediction of gene expression from personal genomes. Genome Biology, 27(1):19, 2026. doi: 10.1186/s13059-025-03926-7.

[16] Brent S. Pedersen and Aaron R. Quinlan. cyvcf2: fast, flexible variant analysis with Python. Bioinformatics, 33(12):1867–1869, 2017. doi: 10.1093/bioinformatics/btx057.

[17] Petr Danecek, James K. Bonfield, Jennifer Liddle, John Marshall, Valeriu Ohan, Martin O. Pollard, Andrew Whitwham, Thomas Keane, Shane A. McCarthy, Robert M. Davies, and Heng Li. Twelve years of SAMtools and BCFtools. GigaScience, 10(2):giab008, 2021. doi: 10.1093/gigascience/giab008.

[18] Žiga Avsec, Roman Kreuzhuber, Johnny Israeli, Nancy Xu, Jun Cheng, Avanti Shrikumar, Abhimanyu Banerjee, Daniel S. Kim, Thorsten Beier, Lara Urban, Anshul Kundaje, Oliver Stegle, and Julien Gagneur. The Kipoi repository accelerates community exchange and reuse of predictive models for genomics. Nature Biotechnology, 37(6):592–600, 2019. doi: 10.1038/s41587-019-0140-0.

[19] Kipoi contributors. kipoiseq: sequence-based data-loaders for Kipoi. https://github.com/kipoi/kipoiseq. Accessed 2026-05-19.

[20] Tinyi Chu, Zhong Wang, Dana Pe’er, and Charles G. Danko. Cell type and gene expression deconvolution with BayesPrism enables Bayesian integrative analysis across bulk and single-cell RNA sequencing in oncology. Nature Cancer, 3(4):505–517, 2022. doi: 10.1038/s43018-022-00356-3.

[21] Mark D. Robinson, Davis J. McCarthy, and Gordon K. Smyth. edger: a Bioconductor package for differential expression analysis of digital gene expression data. Bioinformatics, 26(1):139–140, 2010. doi: 10.1093/bioinformatics/btp616.

[22] Charity W. Law, Yunshun Chen, Wei Shi, and Gordon K. Smyth. voom: Precision weights unlock linear model analysis tools for RNA-seq read counts. Genome Biology, 15(2):R29, 2014. doi: 10.1186/gb-2014-15-2-r29.

[23] Pau Badia-i Mompel, Jesús Vélez Santiago, Jana Braunger, Celina Geiss, Daniel Dimitrov, Sophia Müller-Dott, Petr Taus, Aurelien Dugourd, Christian H. Holland, Ricardo O. Ramirez Flores, and Julio Saez-Rodriguez. decoupleR: ensemble of computational methods to infer biological activities from omics data. Bioinformatics Advances, 2(1):vbac016, 2022. doi: 10.1093/bioadv/vbac016.

[24] Fabian Pedregosa, Gaël Varoquaux, Alexandre Gramfort, Vincent Michel, Bertrand Thirion, Olivier Grisel, Mathieu Blondel, Peter Prettenhofer, Ron Weiss, Vincent Dubourg, Jake Vanderplas, Alexandre Passos, David Cournapeau, Matthieu Brucher, Matthieu Perrot, and Édouard Duchesnay. Scikit-learn: Machine learning in Python. Journal of Machine Learning Research, 12:2825–2830, 2011.

[25] Daniel J. Stekhoven and Peter Bühlmann. Missforest–non-parametric missing value imputation for mixed-type data. Bioinformatics, 28(1):112–118, 2012. doi: 10.1093/bioinformatics/btr597.

[26] Yoav Benjamini and Yosef Hochberg. Controlling the false discovery rate: a practical and powerful approach to multiple testing. Journal of the Royal Statistical Society: Series B, 57 (1):289–300, 1995. doi: 10.1111/j.2517-6161.1995.tb02031.x.

[27] Adam Paszke, Sam Gross, Francisco Massa, Adam Lerer, James Bradbury, Gregory Chanan, Trevor Killeen, Zeming Lin, Natalia Gimelshein, Luca Antiga, Alban Desmaison, Andreas Kopf, Edward Yang, Zachary DeVito, Martin Raison, Alykhan Tejani, Sasank Chilamkurthy, Benoit Steiner, Lu Fang, Junjie Bai, and Soumith Chintala. PyTorch: An imperative style, high-performance deep learning library. In Advances in Neural Information Processing Systems, volume 32, pages 8024–8035, 2019.

[28] Phil Wang. enformer-pytorch: Implementation of enformer in pytorch. https://github.com/lucidrains/enformer-pytorch, 2025. GitHub repository. Accessed 2026-05-18.

[29] Johannes Hingerl. borzoi-pytorch: Pytorch implementation of the borzoi model. https://github.com/johahi/borzoi-pytorch, 2025. GitHub repository. Accessed 2026-05-18.

[30] Johannes Hingerl. johahi/borzoi-replicate-0. https://huggingface.co/johahi/borzoi-replicate-0, 2025. Hugging Face model repository. Accessed 2026-05-18.

[31] The Human Protein Atlas. The single nuclei brain: Explore the expression profiles in single nuclei brain cell types. https://www.proteinatlas.org/humanproteome/single+cell/single+nuclei+brain, 2026. Accessed 2026-05-19.

[32] Andreas Digre and Cecilia Lindskog. The human protein atlas–integrated omics for single cell mapping of the human proteome. Protein Science, 32(2):e4562, 2023. doi: 10.1002/pro.4562.

[33] Kimberly Siletti, Rebecca Hodge, Adriana Mossi Albiach, Kanan Lee, Song-Lin Ding, Lijuan Hu, Bing Zhang, Jaroslav Bendl, Peter Barker, Trygve E. Bakken, et al. Transcriptomic diversity of cell types across the adult human brain. Science, 382(6667):eadd7046, 2023. doi: 10.1126/science.add7046.

